# Phlag: Scalable detection of genomics regions with unexplained phylogenetic heterogeneity

**DOI:** 10.64898/2026.04.10.717778

**Authors:** Ali Osman Berk Şapcı, Shayesteh Arasti, Edward L. Braun, Siavash Mirarab

## Abstract

**Motivation:** Phylogenetic analyses of entire genomes (phylogenomics) have revealed abundant heterogeneity of evolutionary histories. While much has been done to model this heterogeneity and to infer species trees despite it, the current toolkit has a limitation. Most methods assume that gene trees across the genome differ but are all sampled from *the same distribution*, defined by models such as the multi-species coalescent (MSC), and parametrized consistently across the genome. Empirical data strongly suggest this assumption is often violated because the species tree, its parameters, or the process generating the gene trees can all change across the genome. Errors in the data can further compound this heterogeneity.

**Results:** To address this challenge, we define the problem of detecting what segments of the genome are inconsistent with a putative species tree, even after allowing discordance according to MSC. We model gene trees not as a set, but rather as a series (a realization of a stochastic process) along genomic positions. We propose a Hidden Markov Model (HMM) approach applied to quartet statistics measured from gene trees and tie the model to MSC using simulations. The combined use of these three ideas leads to a scalable method called Phlag. On simulated and real data, we show that Phlag can detect many cases of change in underlying evolutionary processes, including reduced recombination rates, population size changes, and admixture, all using the same algorithm.

**Availability and Implementation:** Phlag is available at github.com/bo1929/phlag. All results and scripts can be found at github.com/bo1929/shared.phlag.

## 1. Introduction

A key discovery in phylogenomics has been the observation that the evolutionary history of different parts of the genome exhibits remarkable heterogeneity, and many empirical studies have documented extensive discordance across the tree of life [1]. In response, researchers have introduced models and algorithms that enable species tree inference despite this discordance. Central to these efforts has been a key idea: We can model gene trees as draws from a distribution parameterized by the species tree and designed to capture biological causes of discordance. The multi-species coalescent (MSC) process [2], which models incomplete lineage sorting (ILS), is the most widely used, though gene duplication and loss (GDL), hybridization, and horizontal gene transfer (HGT) have also been successfully modeled [3]. Adopting one of these models, algorithms seek the species tree that best explains the gene tree distribution, essentially attributing all the observed discordance to the modeled biological cause plus random noise. Some methods directly use likelihood under the model [e.g., 4], while others use the model less directly (e.g., the ASTRAL methods rely on the species tree matching the most frequent gene tree for each quartet, as predicted by MSC [5]). Even these less parametric methods often use the predictions of the model directly for *some* steps (e.g., for branch length and support calculation [6]). The reliance on models raises a question: what if the empirical distribution of gene trees does not fit the model?

Several forces can create gene tree distributions that differ from the predictions of the mechanistic models [e.g., 7–13]. The causes of discordance may differ from those assumed by specific models. For example, there may be gene flow or hidden paralogy when the MSC is assumed. Inference of gene trees may introduce systematic errors due to long-branch attraction or inaccuracies in the sequence evolution model [14]. The input data (e.g., the alignment) may contain errors [15]. A subtler challenge is that, even if the assumed model is correct, its parameters, such as recombination rates or population sizes, may vary across the genome. Whether model deviations lead to inaccurate species tree topologies depends on many factors, including the extent of the deviation, but they will clearly affect estimated species tree parameters (e.g., branch lengths and support). Even if the species tree is unscathed, deviations from the model are inherently interesting as they point to changes in the evolutionary process, and masking them reduces our ability to reconstruct a full picture. Thus, instead of asking how much deviations from the model affect the species tree, we ask whether such deviations can be detected systematically.

A key insight from empirical analyses is that model violations can concentrate in gene trees in specific genomic segments [7–11]. The local nature of deviations is helpful in our quest to find model violations because deciding whether a single tree is an independent draw from a (wide) distribution is underpowered. However, once positions are considered, we can look for a stretch of the genome with unusual histories, thereby providing a stronger signal. To formalize this notion, we also need to define what is meant by “unusual”. We assume a species tree is available or can be inferred; then we derive a null gene tree distribution assuming the MSC alone, and look for genomic regions that violate the null hypothesis.

In this paper, we formalize the task of detecting localized deviations from an assumed gene tree model. Given a species tree, focal branches, gene trees ordered by genomic position, and a null gene tree distribution parameterized by the species tree, we seek genomic regions where gene trees do not conform to the null model for the focal branches. The results can be used to further understand causes of discordance and to refine species-tree parameters. We use Hidden Markov Models (HMMs) for this segmentation task and use several techniques to enable scalability, flexibility, and interpretability of results. While our framework is general, we focus on the widely-assumed MSC model. MSC forms a reasonable null distribution because, unlike other causes of discordance, no unusual process is needed for ILS to be present — it is always possible, albeit with probabilities that depend on the species tree. We evaluate the resulting method, called Phlag, through extensive realistic simulations to demonstrate its strengths and limitations. We also show that Phlag can detect known regions with outlier histories in empirical bird and mammal datasets and provide new insights into recalcitrant nodes in the tree of life.

## 2. Methods

### 2.1. Model and problem definitions

Let *T* be a species tree with branch set ℬ, and 𝒳 be the set of all topologies on its leafset. Let [*N*] = {1 … *N*}. We define gene tree sequences {*G*_*i*_ ∈ 𝒳}_*i*∈[*N*]_ and {*A*_*i*_ ∈ 𝒳}_*i*∈[*N*]_. *G* follows coalescent+recombination [16] parameterized by the species tree *T* and numerical parameters *θ*_∅_ such as recombination rates and population sizes. The marginal distribution of *G* follows MSC [2], *G*_*i*_ ∼ 𝒟_∅_(*θ*_∅_). While *G*s are not Markovian, they are often modeled as Markovian, a reasonable approximation adopted for computational expediency [17]. *A* follows an unknown *alternative* process such as gene flow, MSC with parameters different from *θ*_∅_, or unknown distributions resulting from erroneous alignments; its marginal distribution *A*_*i*_ ∼ 𝒟_*a*_(*θ*_*a*_) is unkown but differs from 𝒟_∅_. The alternative 𝒟_*a*_ may be similar to 𝒟_∅_ everywhere except for a subset of focal branches. Let 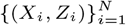 denote a sequence generated by a mixture of these two processes, where *Z*_*i*_ ∈ {0, 1} is a latent indicator variable: *Z*_*i*_ = 0 ⇔ *X*_*i*_ = *G*_*i*_ and *Z*_*i*_ = 1 ⇔ *X*_*i*_ = *A*_*i*_.

We are given a sequence of gene trees *X* = (*X*_1_, …, *X*_*N*_) as a realization of the mixture process (*X*_*i*_, *Z*_*i*_), an unrooted species tree assumed to match the topology of *T*, and a subset of focal branches ℬ′ = {*b*_1_, …, *b*_*m*_} ⊆ ℬ. We have an estimator 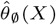 of numerical parameters *θ*_∅_ including branch lengths in coalescent units (CU); note these estimates may have errors. We let 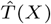 (or 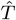 for short) denote *T* together with numerical parameters 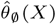. Our goal is to infer the latent sequence *Z* = (*Z*_1_, …, *Z*_*N*_) to distinguish null and alternative gene trees.

### 2.2 Phlag Approach

In principle, we are interested in *P* (*Z* | *X, θ*_∅_, *θ*_*a*_) and thus 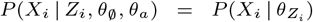. However, computing this likelihood is not practical. First, while we can estimate *θ*_∅_ and thus 𝒟_∅_, we do not know 𝒟_*a*_. Moreover, while *P* (*X*_*i*_ | *θ*_∅_) is, in principle, computable under the MSC given the 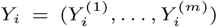, even a viable approximation is not scalable for medium-sized trees [4].

To create a scalable method, we rely on statistics derived from gene trees rather than gene tree likelihood. Let’s assume we have access to a statistic *F* : 𝒳 × ℬ′ → 𝒴 where 𝒴 is some numerical domain (e.g., numbers between 0 and 1). Let *Y* = (*Y*_1_, …, *Y*_*N*_) and *Y*_*i*_ = (*Y*_*i*_^(1)^, …, *Y*_*i*_^(*m*)^) where *Y*_*i*_^(*j*)^ = *F*(*X*_*i*_,*b*_*j*_). Our approach can work with any statistics as long as we can efficiently calculate *F* (*X*_*i*_, *b*_*i*_). We return to *F* later.

Our goal has changed to maximizing *P* (*Z* | *Y, θ*_∅_, *θ*_*a*_). We adopt the Markovian approximation of recombination plus coalescence [17], and we reasonably assume that *Z* is also Markovian. This naturally leads to an HMM, with two states: *null* and *alternative* (Fig. 1B). The HMM framework allows us to *i*) learn HMM parameters (in lieu of *θ*_∅_ and *θ*_*a*_) by maximizing the posterior using a MAP-EM framework, and *ii*) estimate *Z* as the Viterbi path, in lieu of *P* (*Z* | *Y, θ*_∅_, *θ*_*a*_).

**Fig. 1.**
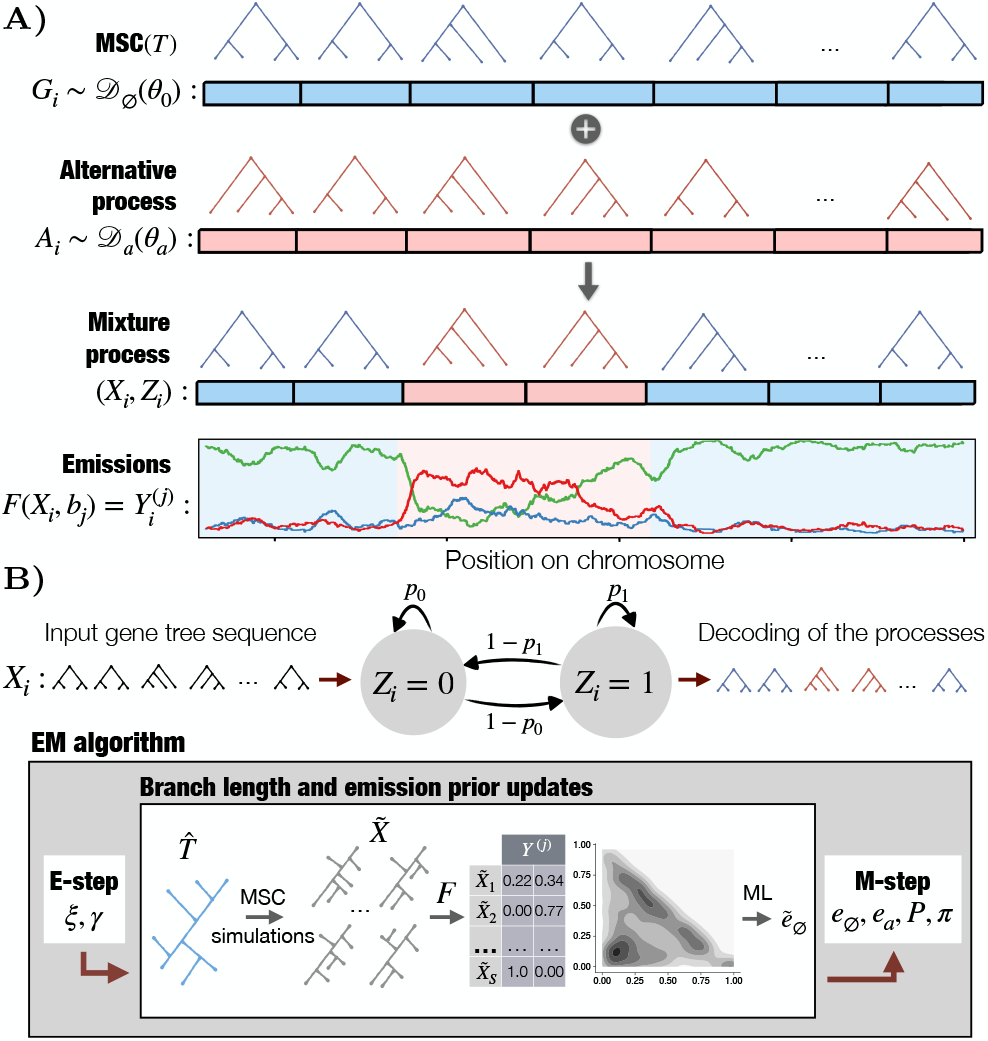
Approach overview. **A)** We model the observed gene tree process as a mixture of gene tree sequences generated by MSC and an alternative unknown process. We represent each gene tree *X*_*i*_ using a statistic *Y*_*i*_^(*j*)^ for each branch *b*_*j*_. **B)** Overview of the inference algorithm. We model *Y* as the output of an HMM with two states, as shown. The EM algorithm for learning HMM parameters is similar to Baum-Welch, except that we introduce MSC-based emission probabilities, 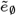, obtained from MSC simulations and used either as the emission distribution for the null state (`MSC-*` modes) or as a Gibbs prior for it (`prior-*` modes). In `*-update` modes, Phlag updates 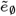 by reestimating the CU branch lengths based on the posterior state probabilities (*γ*) obtained in the E-step.

#### 2.2.1. HMM: Transition matrix

The transition matrix, *P*, is learned from the data, but we designed a prior with two adjustable hyperparameters. Let *β* (default: 4) be the *aprior* expected number of times we transition from null to alternative, and *ρ* (default: 0.9) be the expected portion of the gene trees in the null state. We define a prior on *P* using two Beta distributions:

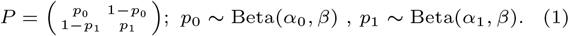

Appendix S2.1 shows that setting *α*_0_ = *Nρ*−*β* and *α*_1_ = *N* (1− *ρ*) − *β* would lead to (*ρ*, 1 − *ρ*) being the expected stationary distribution and *β* transitions from null to alternative. Lower *β* allows us to favor “sticky” [18] states with significantly higher probabilities on the diagonal, while *ρ* controls sensitivity.

#### 2.2.2. HMM: Emission distributions

##### Statistic

The statistic *Y* defined by *F* forms the emissions. A particularly suitable statistic under MSC is the normalized quadripartition quartet score (QQS) [6, 11]. Each internal branch *b*_*j*_ of *T* always has four adjacent branches and defines a quadripartition *A, B*|*C, D*. QQS of *b*_*j*_ for a gene tree *X*_*i*_ (denoted 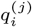) is a number between 0 and 1, showing the proportion of quartets {(*a, b, c, d*) : *a* ∈ *A, b* ∈ *B, c* ∈ *C, d* ∈ *D*} in *X*_*i*_ that have the same topology as *T*. QQS is also defined for NNI rearrangements around *b*_*j*_ (e.g., *A, C*|*B, D* and *A, D*|*B, C*); the three QQS values add up to one, and thus, jointly can be modeled as a simplex, represented as two numbers (the third one is not free). We use these 2-dimensional representations. QQS is suitable because it can be computed in *O*(*n* log^2^(*n*)) with advanced datastructures [19] or in *O*(*n*^2^) with a relatively simple dynamic programming [6], and because quartet trees lack anomaly zones under MSC [5]. Moreover, an estimator 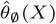 of CU branch lengths of the species tree can be designed solely based on the QQS [6]. We use only this statistic in this paper (using the simpler *O*(*n*^2^) algorithm), leaving the exploration of other options to future work.

While the expected value of QQS under MSC is known, its full distribution has not been analytically derived, and the distribution under the alternative state is unknown. Thus, to parametrize emissions, we approximate their distribution using parametric models 𝒟_*Y*_ parameterized by *e*_∅_ for the null and *e*_*a*_ for the alternative state. Another complexity is that multiple focal branches may be given. To keep the emission space and the number of free parameters manageable, we model branch emissions independently. This allows us to parameterize the emission distribution of each branch separately; thus, we have 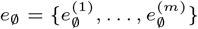 where 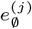 denotes parameters for *b*_*j*_ ∈ ℬ′ (ditto for *e*_*a*_). With this, the emission probabilities are

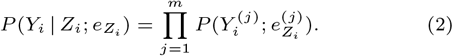

This formulation favors outlier regions that are coincident in all focal branches. Thus, setting ℬ′ to branches that are related to the same outlier region (e.g., neighboring branches, admixing branches) is preferable to an arbitrary subset of branches, which may impact the emission probabilities in unpredictable ways.

We explore several choices for 𝒟_*Y*_, and the framework can easily accommodate others: *i*) We model QQS using a bivariate Gaussian distribution with covariance, leading to 5 free parameters for 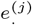. *ii*) To capture the simplex geometry of QQS better, we project QQS values onto Euclidean space via isometric log-ratio (ILR) [20] transform and treat it again as Gaussian. *iii*) We discretize the simplex into three sections, defined by which of the three topologies is dominant. Thus, 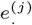 consists of two free parameters. *iv*) (default) We divide the simplex into 6 regions, corresponding to the possible orderings of the QQS values of the three possible quartet topologies, splitting each dominant topology bin into two; this leads to *e*^(*j*)^ having 5 free parameters. *v*) We split the simplex into *k* fixed and equally sized bins in the barycentric coordinate designed for compositional [21] data, leading to only one parameter for the number of bins. *vii*) We use *k*-means to discretize the distribution, leading to *k* − 1 parameters. *vi*) We perform *k*-means in a Euclidean space after an ILR transformation. We decide on the optimal *k* for *v, vi*, and *vii* using the Silhouette score. Due to the independent modeling of emissions, all cases are trivially expanded to *m* > 1 focal branches with a linear increase in the number of parameters.

##### Connecting *e*_∅_ to MSC

For the alternative state, since the distribution of *A*_*i*_ is unknown, we estimate *e*_*a*_ from the data, without informative priors, using the EM algorithm. For the null state, we can use the same approach (`segment` mode), which treats the null and alternative states as two distinct states, without any link to the given 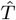 or MSC, effectively segmenting the genome into two alternative signals. To leverage the assumption that *G*_*i*_ should follow MSC, we propose two approaches, each with two flavors, which we will empirically evaluate. Recall we have an estimate 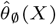 of CU branch lengths [6], which gives us 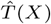. Under MSC, any 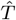 defines a distribution over 𝒳, and we can easily sample that distribution. We simulate a set 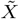 of *S* = 2000 gene trees under MSC using DendroPy [22] and compute 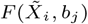 for 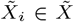 and obtain a distribution over QQS for each branch *b*_*j*_; parameters 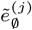 are then trivially fit to the resulting empirical distribution using ML. The least flexible mode (`MSC-fixed`) sets 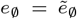, effectively removing *e*_∅_ from EM and leading to zero free parameters for *e*_∅_. An alternative is to make CU lengths free parameters learned by EM. In the `MSC-updated` mode, during EM, we update CU lengths and thus 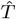(details below) and use the simulations to obtain updated empirical emissions 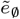.

The two `MSC-*` modes tightly link *e*_∅_ to MSC, which may lead to flagging a region as an outlier even with subtle deviations from MSC. To increase robustness, we can use MSC as a strong prior while letting EM learn *e*_∅_ from the data. This informative prior can be fixed (`prior-fixed`) or updated (`prior-updated`) as EM progresses, corresponding to the empirical Bayes approach. The prior is defined independently per branch. Consider the empirical emission 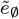 obtained under MSC using simulations. Instead of fixing *e*_∅_ to 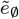, we can use it to define a distance-based Gibbs prior on the space of probability distributions:

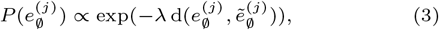

where 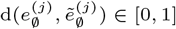 is the distance between the emission distributions parameterized by 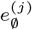 and 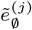, and *λ* is a hyperparameter controlling the strength of the MSC-based prior. Implementing this prior simply requires adding the Hellinger distance to the reference empirical MSC distribution 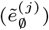 as a regularization term to the log likelihood, pushing the learned posterior towards the baseline. Increasing *λ* results in relying on MSC more, and setting it to 0 reduces us back to the `segment` mode. We use the Hellinger because unlike the KL divergence, it is symmetric, bounded, and always defined.

### 2.3. Parameter inference using EM decoding

The overall inference procedure is summarized in Algorithm 1. We infer the parameters of our HMM using an extension of the Baum–Welch (BW) algorithm, with extra steps for updating the empirical distributions 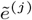. After training parameters using EM, we use the inferred parameters (*e*_∅_, *e*_*a*_, *P, π*) and decode the Viterbi path for *Z* (ViterbiDecoding), producing the decoded null versus alternative inference.

#### Algorithm 1

Inference of parameters and decoding. Initialize initailizes parameters. Each EM iteration requires *O*(*mN*). EstimateBranchCU updates CU branch lengths using Equation (4) in *O*(*nN*). Each update of 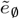 is *O*(*mnS*).

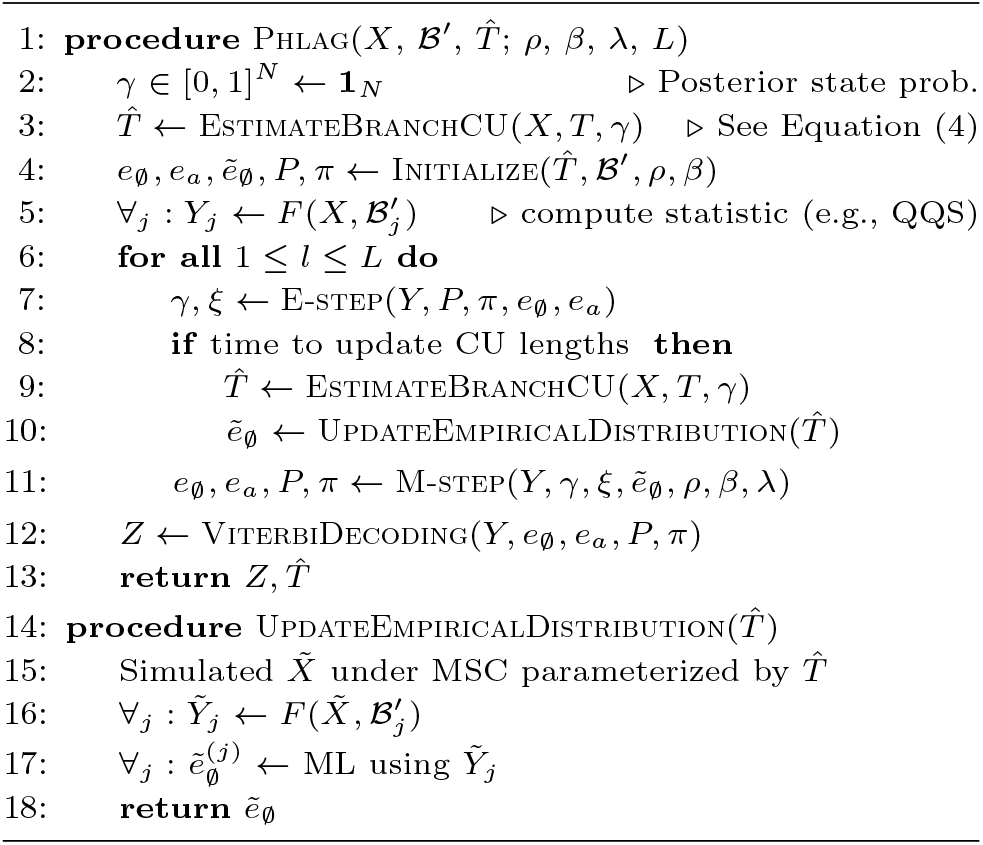

In each iteration, E-step computes the posterior state probabilities *γ*_*i*_ and pairwise transition probabilities *ξ*_*i,i*+1_ using the standard forward–backward algorithm. In the M-step, we update the emission parameters *e*_∅_ and *e*_*a*_, the transition matrix *P*, and the initial distribution *π* by maximizing the expected complete-data log-posterior and the conjugate Beta priors on *P*. The initial transition matrix *P* is sampled from its conjugate Beta prior, while the initial state distribution *π* and the alternative emission parameters 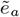 are initialized randomly. We always initialize *e*_∅_ to the simulated 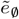, except for the `segment` mode. In the `segment` mode, these steps all follow the traditional BW algorithm. Implementing `MSC-fixed` and `prior-fixed` modes are also straightforward: 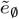 is precalculated using 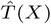 before BW; in the first case, emissions of the null state are fixed and removed from BW; in the second case, prior *P* (*e*_∅_) is added as a regularization term (see Appendix S2.2).

In the two `*-updated` modes, we recalculate the CU branch lengths in *O*(*nN*) and redo simulations to update 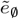 in *O*(*mnS*) [6]. To save computation, we follow an irregular regime to update more frequently in the early iterations. By default, we update 5 times, first at the 25th iteration, and doubling the wait time after each update. Note that from the E-step, we have the state posterior *γ* = (*γ*_1_, …, *γ*_*N*_), each *γ*_*i*_ = *P* (*Z*_*i*_ = 0 | *X, θ*_∅_, *θ*_*a*_) giving the probability of each gene tree being in the null state according to current parameters. We use these to reestimate the branch lengths of 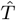 via a weighted quartet-based estimator approach that extends the MAP estimator of Sayyari *et al*. [6] (EstimateBranchCU) to incorporate the posterior *γ*_*i*_:

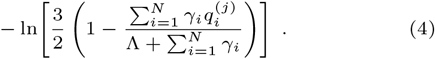

where Λ = 1 is the prior on speciation rate according to the Yule model [6]. Given the updated branch lengths, we resimulate gene trees to update the prior parameters 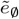.

## 3. Experimental setup

### 3.1. Simulation studies

We first evaluate Phlag modes on simulated gene tree sequences, with the alternative state changing either population size or recombination rate (E1). We then extend the simulations to include admixture and benchmark Phlag, using the default choice of *𝒟*_*Y*_ (topology order) in the `prior-updated` mode, against alternative approaches (E2). While no other dedicated tool exists for the exact task studied here, we can compare with some baselines. *i*) PhylteR [23] (v0.9.12) treats gene trees independently (i.e., is not position-aware) and detects deviations from overall gene tree distribution using gene-specific distance matrices, without relying on a species tree. We evaluate different settings of the sensitivity hyperparameter *k* ∈ {0.2, 0.5, 1.55, 3.00} and report results at multiple levels. *ii*) Bipartition+HMM is similar to Phlag but uses binary emissions based on the presence/absence of focal branch bipartitions in each gene tree, using neither priors nor the MSC. *iii*) Moving average (MA) computes the MA of the QQS of the dominant topology [11] and use it to compute a *z*-score defined as 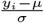, where *µ* and *σ* are the mean and standard deviation, and *y*_*i*_ is the average of QQS on the window centered at position *i*. We use a window size of 100 and mark a position as an outlier if its *z*-score > *u*. We test for *u* ∈ {0.1, 0.2, 0.3, 0.5, 1}.

In all simulation experiments, we apply Phlag and bipartition+HMM on a single focal branch of interest if the estimated branch length is less than 1 CU; for longer branches where QQS is often 0 or 1, we add adjacent internal branches to *B*′ to increase the signal. We keep Phlag’s hyperparameters fixed across different scenarios: *β* = 4, *ρ* = 0.9, and *λ* = 1.5. Since other methods are not aware of the null model to distinguish between the two binary states (e.g., bipartition HMM and `segment` mode), we assign the majority-predicted state to 0 and compute evaluation metrics accordingly.

We use `msprime` [24] to simulate genealogies under the Hudson coalescent model, following recent work [12]. As our null demographic model, we choose the avian phylogeny inferred by Stiller *et al*. [25], which consists of 363 taxa. We configure model parameters using empirical branch length estimates in three units: time, CU, and substitution units (SU). Each population’s effective size (2*N*_*e*_) is set to *t*/(*G* · CU), where *t* is the length in time, *G* is the generation time set to 10, chosen as an approximate midpoint in compilations of avian and mammalian generation times [26, 27]. We simulate 6Mbp alignments under the GTR model with rate heterogeneity across the genome, avoiding unrealistic strict molecular clock (i.e., ultrametricity) by incorporating realistic rate multipliers, obtained from empirical ^*SU*^/*t* rates. From these alignments, we estimate gene trees for *N* = 1500 uniformly spaced 500bp loci using IQ-TREE [28], skipping 8Kbp after each locus. See Appendix S2.3 for details. The discordance between the resulting gene trees and the species tree closely matches that observed for the real Stiller *et al*. dataset (Fig. S1).

We create replicates by combining gene trees simulated under the null model and those from an alternative condition. We randomly select a starting position and substitute a subsequence of gene trees with those simulated under the alternative process. The portion of alternative gene trees is set to {2%, 5%, 10%, 15%, 20%, 25%}. For the alternative region, we keep all parameters and the demographic model unchanged, except for a single branch, on which we introduce a substantial change to a single model parameter. For 40 randomly selected branches, we either increase or decrease the population size by 10×, resulting in 20 cases in each direction. For recombination rate changes, we increase the rate 10× for 25 branches and suppress recombination by decreasing the rate 1000x for another 25 branches. For ancient admixture, we simulate pulse migration events between two populations for 10 branches (6 events occurring 16–26 Mya and 4 events occurring 60–70 Mya). We set the gene flow proportion in the outlier region to 90% to create conditions similar to selective introgression, where the admixture signature is strong in one region [10]. In total, we performed 600 replicate simulations.

### 3.2. Biological data

We tested Phlag on two large empirical phylogenies (E3: avian and E4: mammalian). We selected these because each has genome-wide alignments and known examples of outlier regions among internal branches [11, 29]. For Phlag analyses, we keep *λ* the same as in the simulations but increase *β* to 100, thereby maintaining a similar expected number of transitions per gene tree (e.g., 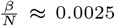 for the avian dataset), and *ρ* to 0.95. This allows for more fine-grained detection of outliers along longer gene tree sequences (up to 40,000). Phlag is applied to one focal branch at a time, without including neighbors.

For birds, we study 39,849 gene trees sampled from five macrochromosomes from a 363-taxon dataset [25], selecting 1Kbp subalignments with minimum missing data from each 10Kbp segment, aiming to break dependencies across gene trees to better comply with MSC. We focused on 20 branches, selecting from 33 basal branches (60-68 Mya) highlighted by Stiller et al. [25] that had quartet support below 0.5 and were within the Neoaves. For mammals, we used the alignment by Foley et al. [30] and focused on chromosome 3 because there is a well-characterized unsual ILS region in humans and chimpanzees located at band 3p21.3 [31]. We applied Phlag on all branches under key orders: Carnivora, Chiroptera, Primates, Artiodactyla, and Rodentia. Among these, we only retained the ones that define a quadripartition in at least 90% of the gene trees, resulting in 136 branches. Since no gene trees were available, we inferred 19,465 gene trees using a protocol similar to Stiller et al. [25], with the same segment and subalignment sizes, and running IQ-TREE [28] under the `GTR+G4` model with approximate Bayesian support. To identify the positions of these loci on non-human genomes, we matched the names of coding genes in GTF files downloaded from Ensembl. We compared the annotations for the human GRCh38.115 assembly with a single high-quality assembly from each clade (bat mRhiFer1 v1.p.115, cow ARS-UCD2.0.115, dog ROS Cfam 1.0.115, and mouse GRCm39.115).

## 4. Results

### 4.1. E1: Comparing Phlag vairants

We first ask if expanding ℬ′ by adding the neighbor branches of a focal branch (where the MSC is violated) improves results. Answers depend on the CU length estimate of the focal branch and the type of change. For changes in recombination, expansion dramatically helps recall, particularly for long branches; for population size changes, expansion can inflate the false positive rate (FPR), especially for short branches (Fig. S2). We adopt expanding when the estimated focal branch length is above 1 CU as the default strategy.

The choice of emission type has a substantial impact on both recall and precision, regardless of the scenario. Among the evaluated options, using a Gaussian distribution performs the worst (Fig. 2). Projection onto Euclidean space via ILR helps recall at the expense of the FPR. This result can be explained by QQS distributions being often multimodal, particularly near terminal branches. All the discretized metrics perform better. Although 3-bin dominant-topology emissions achieves the highest recall, it suffers from high FPR and is clearly less specific than that of the 6-bin topology order. Discretization using *k*-means, with or without ILR projection, consistently yields lower recall and higher FPR than the simpler topology-order representation and typically requires more parameters (i.e., *k* > 5). Compared to barycentric bins, which tend to achieve the lowest FPR among the discrete emissions, topology order improves recall by 28% with only an 8% increase in FPR. Together, these results motivate the 6-bin 5-parameter topology order as a robust emission type, used hereafter.

**Fig. 2.**
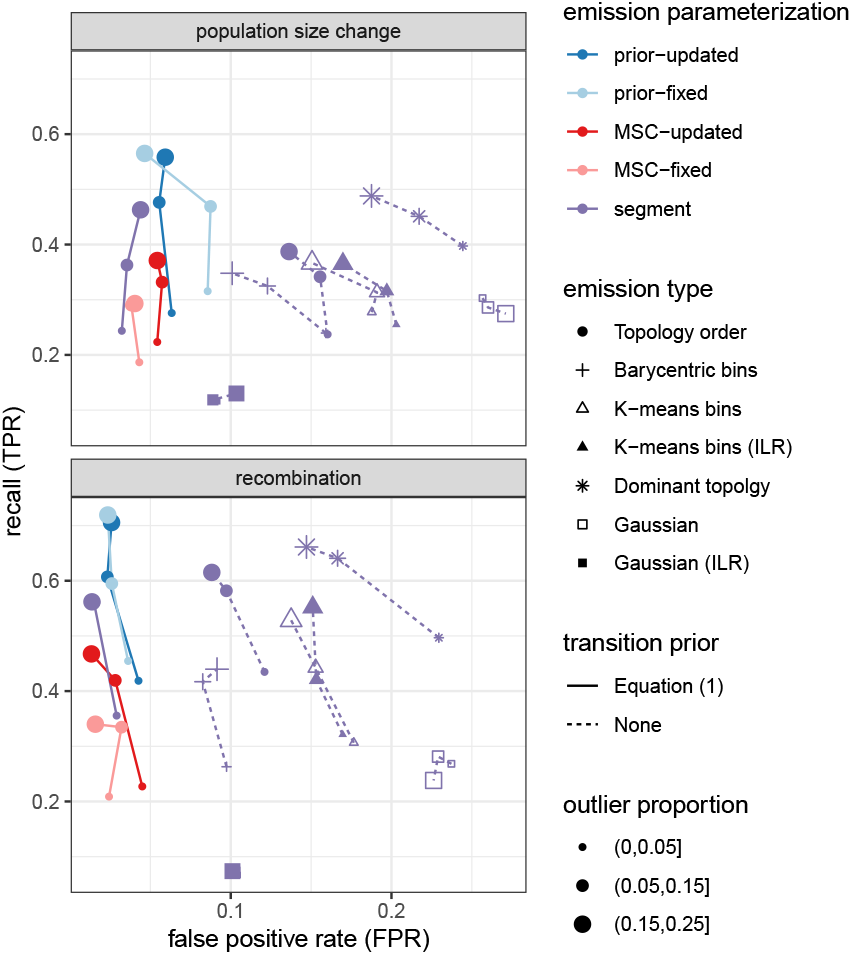
Comparing Phlag variants, differing in emission type, emission parameterization, and whether they put a prior on the transition matrix. Recall and FPR are defined as ^*T P*^/(*T P* + *F N*) and ^*F P*^/(*T N* + *F P*), resp.

Enabling transition priors by Eq. (1) provides a dramatic reduction in FPR, with a slight decrease in recall observed in recombination scenarios and a slight increase for population size (Fig. 2). These improvements are particularly pronounced at higher outlier proportions, with an overall 73% decrease in FPR and a 5% increase in recall, leading us to adopt the feature.

Various modes for tying the null emissions to the MSC model differ in performance. The purely data-driven `segment` mode outperforms both `MSC-fixed` and `MSC-updated`, consistently achieving higher recall and comparable or better FPR across both scenarios. Results suggest that tightly coupling *e*_∅_ to MSC can be disadvantageous, potentially due to inaccuracies in branch length estimation or sensitivity to small deviations from MSC. In contrast, the `prior-updated` and `prior-fixed` modes substantially improve recall compared to data-driven `segment` at only a slight cost to FPR. Finally, the empirical Bayes approach of updating the prior (`prior-updated`) leads to small but noticeable improvements in FPR compared to a fixed prior for population size changes. Thus, we use it as the default.

### 4.2. E2: Benchmarking Phlag against other methods

We compare Phlag against the best-performing configuration of three methods (PhylteR with *k* = 1.55, MA with *u* = 0.3, and bipartition-HMM with CU-based branch extension, identical to Phlag; see Fig. S2). Phlag is the only method that outperforms or is competitive with other methods across all conditions (Fig. 3a); the alternative methods are each better than Phlag under some conditions, but fully fail under others. Also, only Phlag achieves near-perfect accuracy under some conditions.

**Fig. 3.**
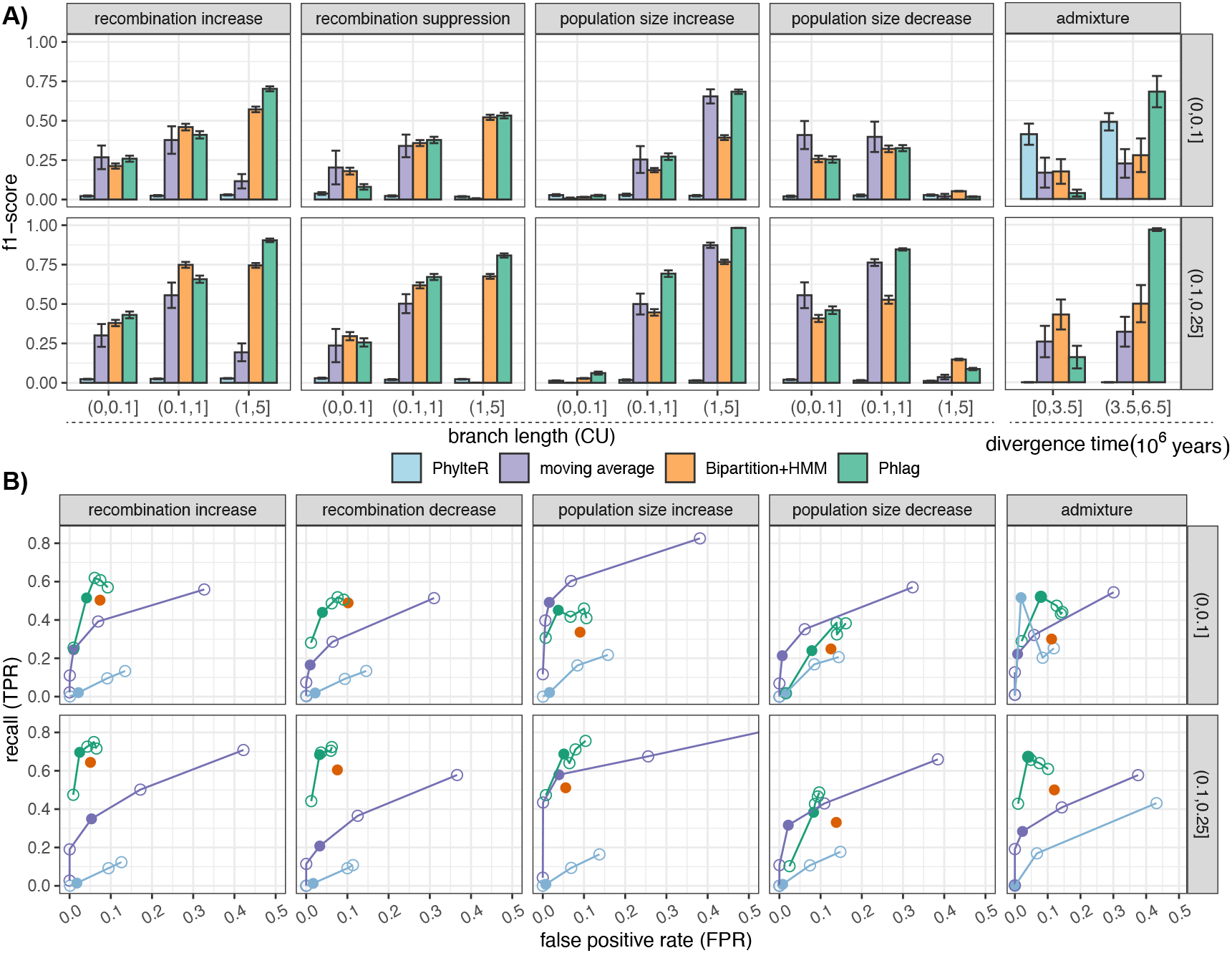
Phlag versus alternative methods on simulated gene trees. **A)** Showing f1-score across different types of outliers (box columns), CU branch lengths where the change occurs (*x*-axis), and total portion of the gene trees affected (box rows). For admixture, *x*-axis shows the divergence time in million years between admixing populations. **B)** Showing TPR and FPR as detection hyperparameters change. For Phlag, we vary *ρ* ∈ 0.99, 0.9, 0.8, 0.7, 0.5 in Eq. (1) (see Fig. S3 for *β*). For Phylter and the moving average, we adjust *k* and *u*, respectively. Solid dots correspond to settings shown in A).

The position-agnostic PhylteR fails to identify changes in recombination rates and population sizes (Fig. 3a). However, it performs well for admixture detection when the introgressed region is 10% or less. When divergence between the admixed populations is short (<3.5My), Phylter is the best method. Its performance degrades substantially as the proportion of admixed gene trees increases. Phlag outperforms alternatives for admixture detection at higher divergence times, achieving over 90% f1-score when outlier proportion is 10–25%. The alternative HMM-based approach, bipartition-HMM, performs well for changes in recombination rate but is substantially less accurate for population size increases. Even for recombination rate change, Phlag performs better than bipartition-HMM on longer branches (CU>1), achieving f1-scores of >80% when affected gene trees constitute at least 10% of the genome; for shorter branches (CU≤1), both methods perform comparably. Moving-average (MA) performs well for population size detection, matching, or slightly suppressing Phlag in several cases. However, it fails to detect recombination changes, especially for changes occurring on branches with CU>1.

Not all types of change are equally challenging. For example, the impact of population size increase or decrease depends on branch length. All methods fail to detect an increase on a short branch (CU≤ 0.1), as background ILS is already high and masks further ILS increases. As branches get longer, the f1-score of all methods improves, with Phlag achieving the best performance. Conversely, population size decreases (reduced ILS) are difficult to capture on long branches where ILS is low to begin with. Most methods, and Phlag in particular, tend to work better when outliers are 10–25% rather than <10%.

Phlag, MA, and Phylter have hyperparameters that can trade off FPR and recall. For instance, the default *k* value of PhylteR is 3.0, but 1.55 is used as an alternative for high-sensitivity analysis [23], which also improves recall for admixture detection in our analysis. The effect of changing *u* in MA is more clear (Fig. 3b); *u* = 0.3 yields the highest f1-score, but increasing it further worsens FPR rapidly, with little gain in recall. In contrast, Phlag is robust to changes in *ρ*, except for at extremes (e.g., *ρ* = 0.99). Increasing *ρ* beyond 0.90 improves recall in many cases but not all. Nevertheless, in recombination suppression, Phlag maintains FPR below 0.1 across all tested *ρ* values, retaining relatively high recall (above 0.6 and 0.4 for outlier proportions in (0, 0.1] and (0.1, 0.25], respectively) for *ρ* < 0.99. At comparable FPR levels, Phlag consistently outperforms alternative methods for recombination rate and population size changes for portions > 0.1. Similarly, changing *β* impacts results only mildly (Fig. S3).

### 4.3. E3: Applying Phlag to Avian phylogeny

We use two criteria to evaluate Phlag on biological data: *i*) its ability to identify regions known to have unusual gene tree distributions, and *ii*) whether it can flag branches that have been difficult to resolve in earlier studies. Phlag was very successful based on the first criterion; it identified a region of chromosome 4 known to have an unsual signal of ILS (Fig. 4). Since the deviation in the gene tree distribution in this region is extremely strong [11], the original study detected it by visual examination, making it reassuring that the signal is detected by Phlag. In fact, judged by Hellinger distance, this known outlier was the strongest example of violating MSC across the tree. Consistent with the “Columbea” rearrangement (Fig. 4) in this region, four adjacent branches are affected by the anomaly. Note that this region has been implicated as a major difficulty in resolving the Neoavean first divergence [11, 25].

**Fig. 4.**
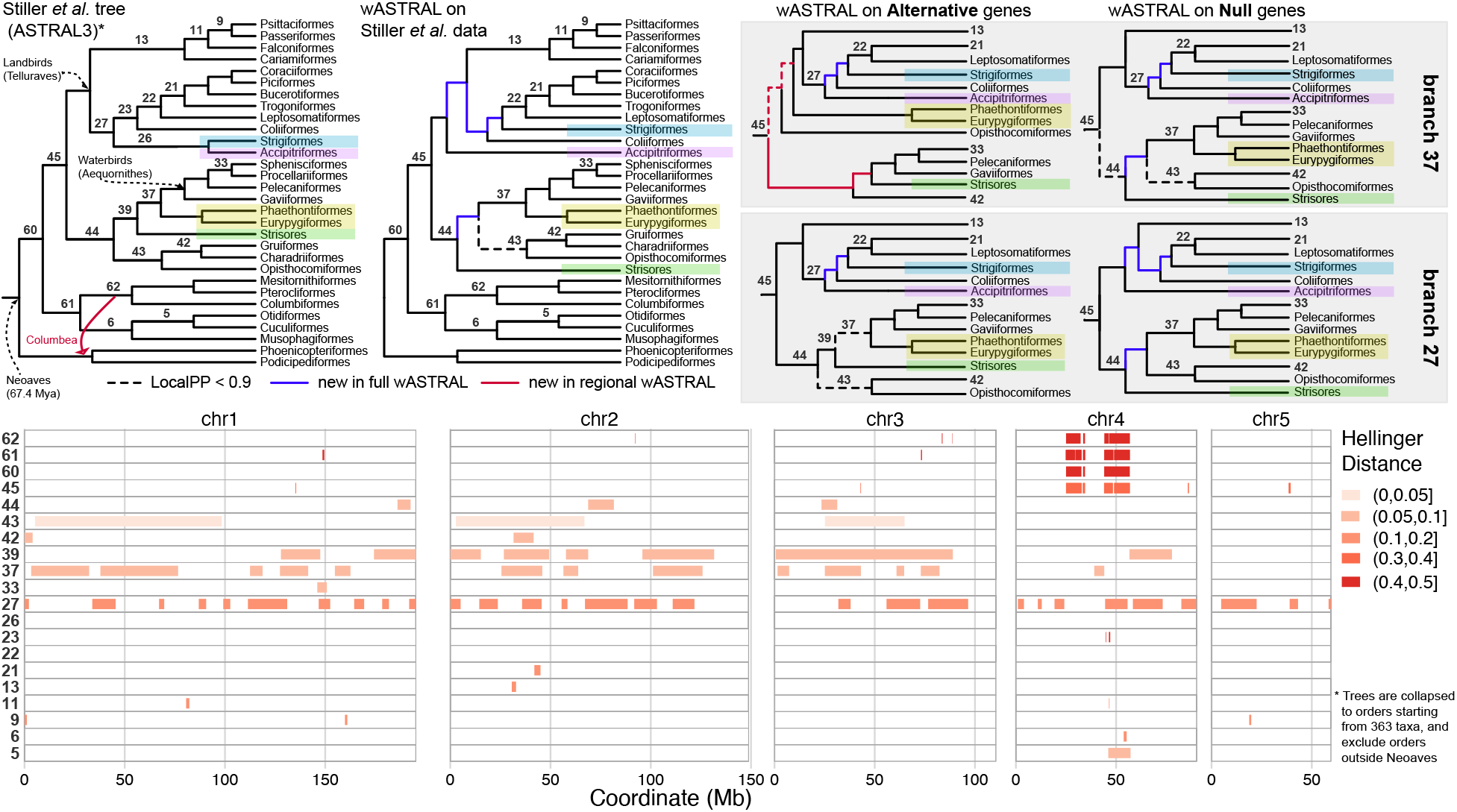
Phlag analysis of bird macrochromosomes. Bottom panel shows Phlag-identified regions, colored by Hellinger distance between the emission distributions of the states, for 20 branches near the base of Neoaves and with QQS < 0.5. Branches are labelled on the Stiller et al. [25] tree (top left). The four highlighted groups have been unstable in earlier analyses [32]; running wASTRAL instead of ASTRAL3 changes four (blue) branches. Removing or only using loci from flagged regions for corresponding branches 27 and 37 leads to wASTRAL trees that reveal hidden support for alternative positions of these four taxa (see Fig. S5), consistent with hybridization. Known [11] strong outliers in Chr 4 lead to support for Columbea, shown on the left.

Phlag was also able to identify the Z chromosome (the sex chromosome heterogametic in female birds) as a region with an unusual gene tree distribution Fig. S4. This was expected because the Z chromosome, by theory, has a smaller effective population size than autosomes and a higher rate of sequence evolution due to the “fast-Z” effect [33].

We next turn to the second criterion. Among the reasons that branches might have been difficult to resolve, reticulate evolution is likely to result in spatial clustering of genes with similar ancestry. Phlag revealed many segments with unexpected signal on five of the branches we examined (branches 27, 37, 39, 43, and 45; Fig. 4), all of which have been difficult to resolve in earlier studies [32], including by genome-wide data [25]. In fact, analyses of Stiller et al. [25] were inconclusive about the placement of Phaethontimorphae and Strisores (branches 37 and 39), and that of Accipitriformes and Strigiformes (branches 26 and 27), and they suggested that these might have a reticulate origin. Our results are consistent with a reticulate origin. Phlag identified large blocks with gene tree distributions that deviate from the null hypothesis for branches 27, 37, and 39. As expected for reticulate evolution, the wASTRAL tree based on gene trees from the outlier regions moved these clades (Figs. 4 and S5). For branch 37, Phaethontimorphae becomes sister to landbirds with outliers, while it remains with waterbirds with the null gene trees.

Patterns for branch 27, which unites the putative clade Afroaves (Fig. 4), were more complex because it is so unstable that a wASTRAL reanalysis of the Stiller et al. [25] gene trees fails to recover it. Instead, it puts Accipitriformes sister to all landbirds, a pattern observed before [25, 32]. wASTRAL analysis of alternative state gene trees moved Accipitriformes to a clade including Strigiformes and several other orders, whereas analysis of null trees recovered the same position as the full wASTRAL tree. These results are consistent with one or more reticulation events at the base of landbirds.

### 4.4. E4: Applying Phlag to Mammalian phylogeny

The known unsual ILS region in humans and chimpanzees, located at band 3p21.3 [31] was evident in the Phlag analysis (Fig. 5). The Hellinger distance from the null for this region was not particularly high and it was fragmented into several outlier segments; this probably reflects the variation in the degree of ILS in this region evident in previous analyses [29]. The chromosome numbers for the human 3p21.3 genes differ in other clades (Fig. 5), but a set [34] of eight putative tumor-suppressor genes in this region have fully conserved gene order in the four genome assemblies we examined. Phlag identified outlier segments that overlap with that region in several clades (Chiroptera, Yangochiroptera, Cercopithecinae, and Sciuromorpha). CASTER also revealed a non-ILS signal in this region across many clades [11], and introgression has been proposed for Cercopithecinae [35]. The outliers in Chiroptera show patterns similar to Homininae, with modest QQS asymmetry (Fig. S9), while QQS asymmetry is much larger in Rodentia, in one case locally exceeding the QQS of the main tree (I161.Rodentia; Fig. S9), suggesting introgression.

**Fig. 5.**
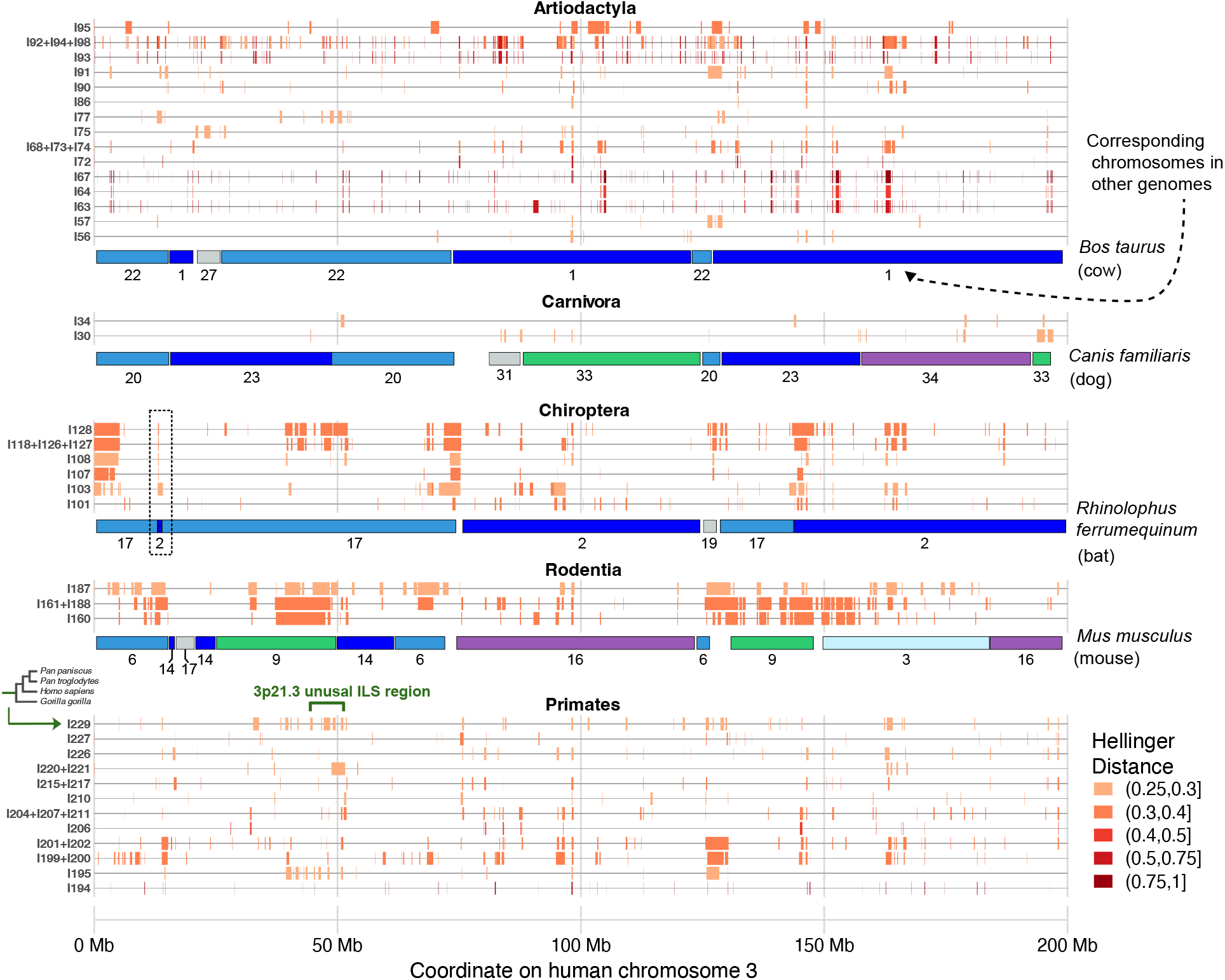
Outlier regions with Hellinger distance >0.25, detected by Phlag in selected mammals for genes located on human chromosome 3. Lines correspond to branches in the mammal tree and are identified using edge labels (see Fig. S6). Clades are collapsed into a single row if the taxonomic common ancestor of all taxa under them is the same. Colored boxes below each of the non-primate clades correspond to chromosomal blocks in selected high-quality genome assemblies for each clade. The box with dashed lines indicates a region with a small rearrangement. Chromosomes with a small number of human chromosome 3 loci or rearrangements within the blocks are not shown. The branches shown in this figure contain outlier regions with high distances; see Fig. S7 for other branches, and Fig. S8 for *β* = 50 (resulting in a value of ^*β*^/*N* similar to that of the avian dataset). See also Fig. S9.

Beyond the known region, as expected, most branches had few outliers (Fig. S7). The large number of chromosomal rearrangements in mammals [36] can complicate Phlag analyses by breaking up contiguous regions. However, some outlier regions coincide with rearrangements. One such case is a short segment located on chromosome 2 of *Rhinolophus ferrumequinum* that is an outlier for the *Rhinolophus* branch (Fig. 5; dashed box). This region includes four coding genes, *IQSEC1, NUP210, FBLN2*, and *WNT7A*, one of which, *WNT7A*, was among the set of rapidly evolving genes in an artiodactyl genome [37]. Artiodactyls showed many outliers, with some cases in Bovidae having high Hellinger distances from the null. Extensive introgression among cattle is documented using identity by descent (IBD) [38], though previous work has mostly identified the much stronger signal on chromosomes 18 and 28; Phlag may be able to find less obvious signals from gene trees. Many branches in mammalian phylogeny are characterized by introgression [30, 39], making the extensive output by Phlag realistic and emphasizing the potential for Phlag to identify targets for molecular evolution studies.

## 5. Discussion and Conclusions

We introduced Phlag, a method for segmenting the genome into regions that do or do not follow expectations of MSC for given branches of a species tree. By doing so, Phlag helps elucidate the complexity of evolutionary history; it can be used to improve the species tree by removing outlier regions or implicate species network when alternative states are prevalent. Our HMM-based framework offers much flexibility, which we have only started to explore. Our results confirm that priors for transitions, which encourage contiguity, and for emissions, nudging the null state toward MSC, were essential to high accuracy. The priors introduce three hyperparameters; users are encouraged to explore their impact on their real data.

Our results also showed that the statistic used by HMM as emissions matters, with different discreteizations of quartet scores (QQS) leading to large differences in accuracy. Besides QQS, many other forms of signal can be incorporated, starting with the presence of bipartitions, which showed some promise in our data. Even more advanced metrics, specifically designed to interrogate positional gene tree signal, are available in the literature, including TWISST [40] and SplitScores [41]. We note that these metrics can be combined with Phlag as the statistical function *F* that defines emissions. Future work should explore such combinations and whether the optimal choice depends on phylogenetic depth (e.g., recent or old admixture events).

Two limitations of the current Phlag method need caution. First, it has only two states. In real data, the non-MSC signal can vary (e.g., suppressed recombination in one region and alignment error in another). It is not immediately clear how the model would perform in the presence of multiple non-MSC signal with highly different emissions, and whether a single alternative state can be sufficient. Future research should extend Phlag to allow multiple alternatives, which is technically easy, but requires care in terms of over-parameterization and appropriate priors. An alternative is an iterative application: run Phlag on the full dataset, remove strong outliers, and rerun. In fact, we did just that on the bird dataset; our initial analyses that included the Z chromosome identified it as an outlier, reducing its ability to flag autosomal outlier regions (Fig. S4). We then removed chromosome Z and found other outliers. Second, Phlag requires gene tree orders but synteny breaks in deep time, and orders change across the tree. Future work can explore applying Phlag to each branch based on the orders for that branch, with ancestral reconstruction of gene orders. The per-branch nature of Phlag enables such analyses.

## Supplementary Material

**Fig. S1.**
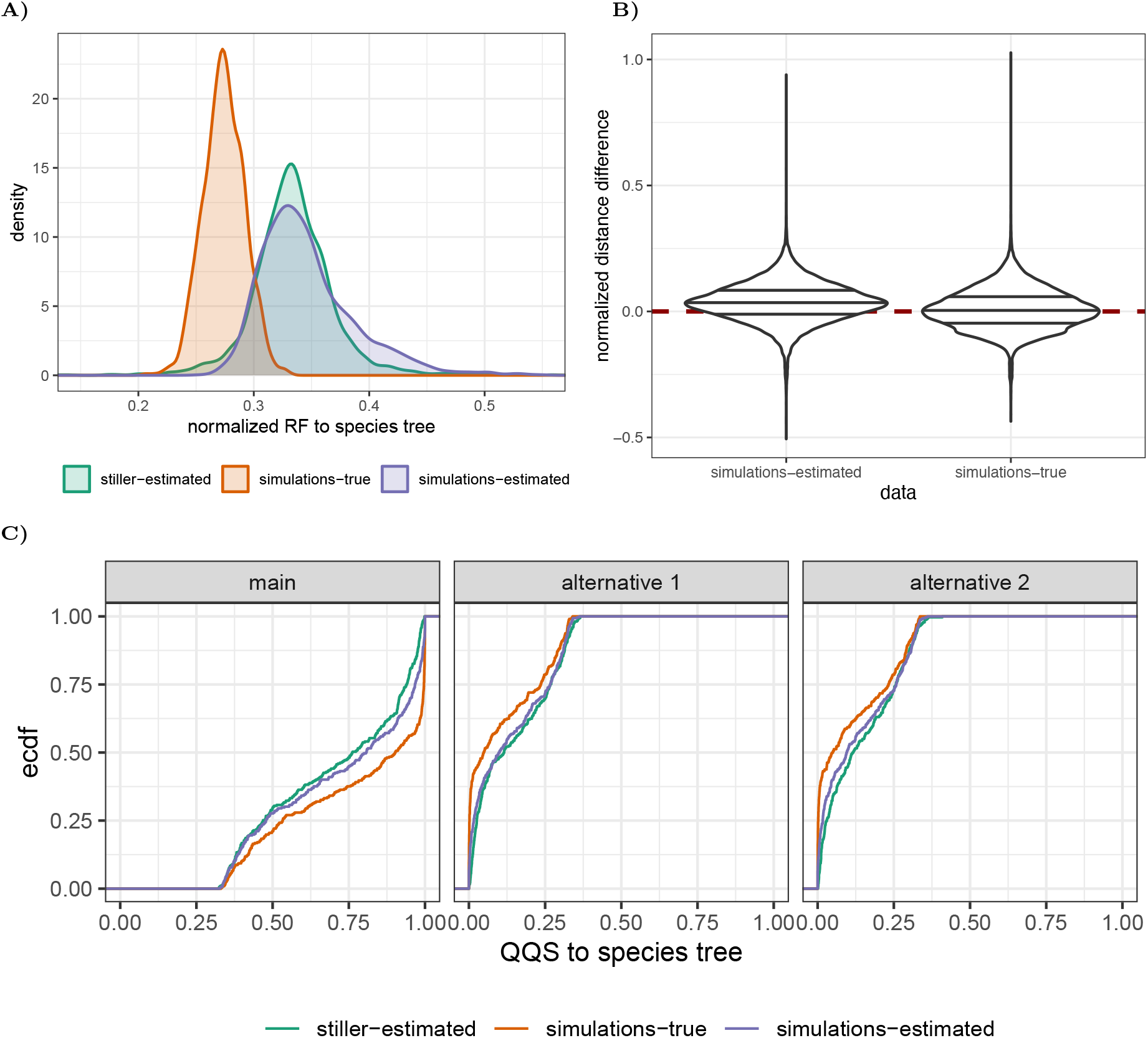
Summary statistics of the simulated gene trees. We show three sets of gene trees: *i*) stiller-estimated: gene trees estimated by Stiller et al. [25] from real data; *ii*) simulations-true: “true” simulated genealogies generated using msprime; and *iii*) simulations-estimated: gene trees estimated using IQ-TREE from simulated sequences of 500 bp regions. A) Distribution of normalized RF distances between gene trees and the reference species tree. Note the close match between the simulations-estimated gene trees used in our analyses and the stiller-estimated trees. B) The difference in pairwise distances (in substitution units) between simulated and Stiller et al. [25] gene trees. We compare the average pairwise distance between a pair of taxa across all gene trees for simulations 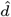 versus Stiller et al. [25] gene trees *d*^∗^. The distributions are shown over all pairs of taxa. We show 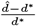. Distributions are centered around zero, showing that selected substitution rates lead to gene trees that match empirical data. C) Empirical distribution function of Quadripartition Quartet Scores (QQS) of gene trees with respect to the reference species tree. Note the close match between the empirical data (stiller-estimated) and the simulated data (simulations-estimated).

**Fig. S2.**
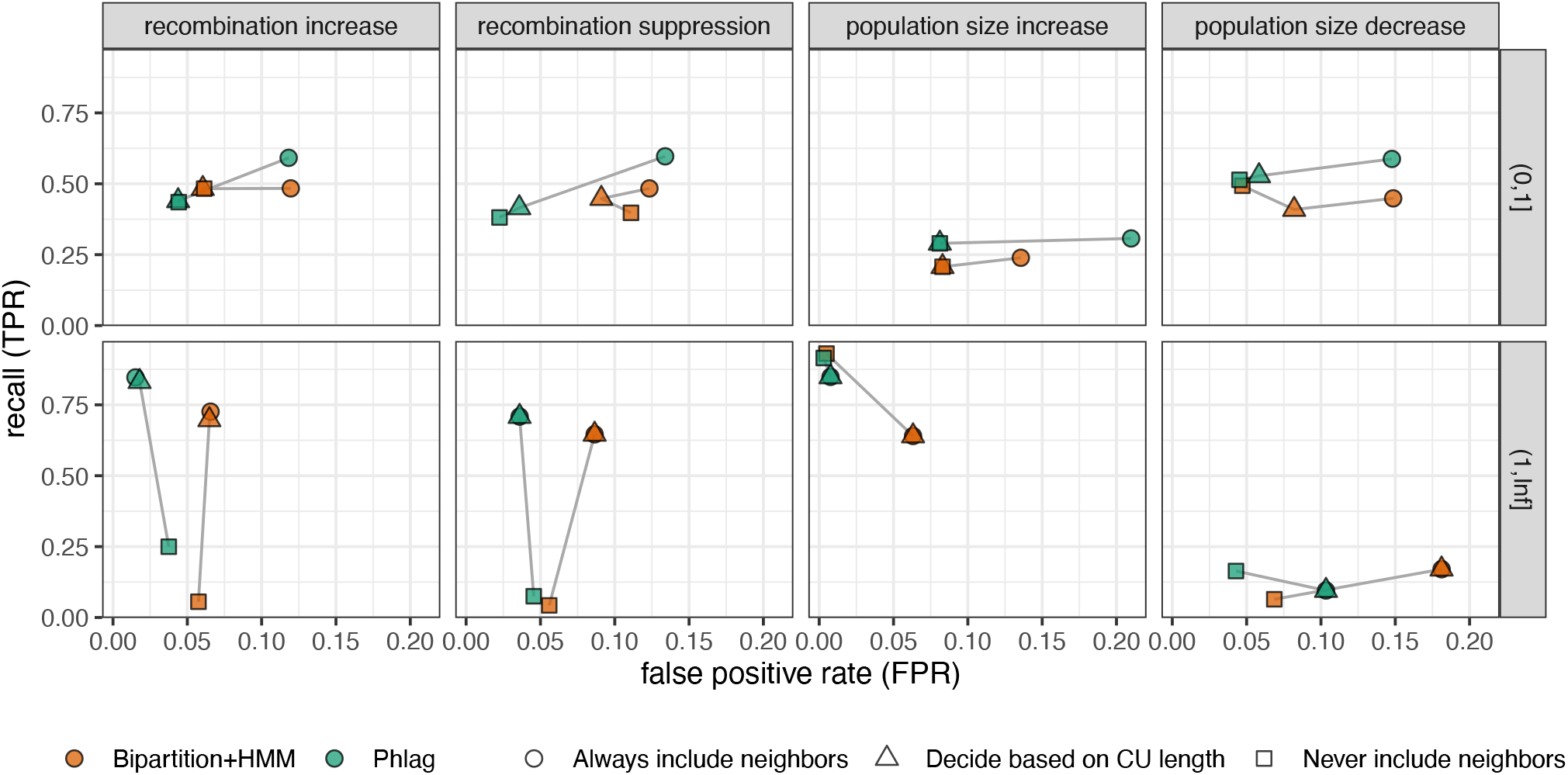
Demonstrating the impact of expanding ℬ′ by adding neighbor branches of the focal branch. We show the false positive rate (FPR) on the *x*-axis, and the true positive rate on the *y*-axis. Shapes indicate the expansion strategy. “Decide based on CU length” strategy adds neighbor branches if the focal branch length estimate is greater than 1 CU. We note that the decision is made based on the estimated branch lengths. We show the true CU branch lenghts in panel rows.

**Fig. S3.**
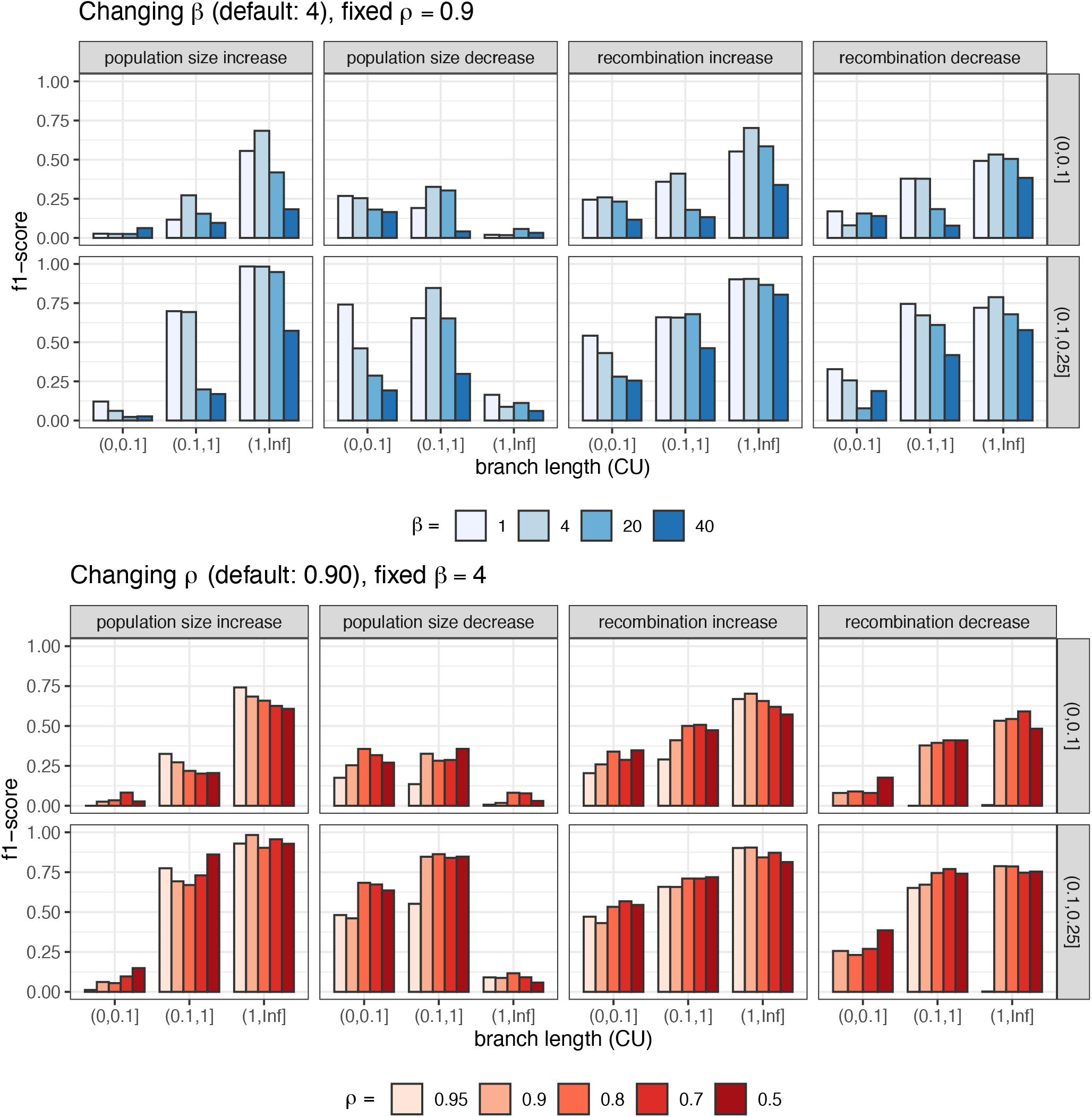
Showing the impact of varying pior hyperparameters *β* ∈ {1, 4, 20, 40} (top) and *ρ* ∈ {0.50, 0.70, 0.80, 0.90, 0.95} (bottom) defined in Eq. (1) on the performance shown as the f1-score on the *y*-axis. Branches are binned based on CU lengths on the *x*-axis. Panel columns are for different types of alternative states, and panel rows correspond to the portion of affected gene trees.

**Fig. S4.**
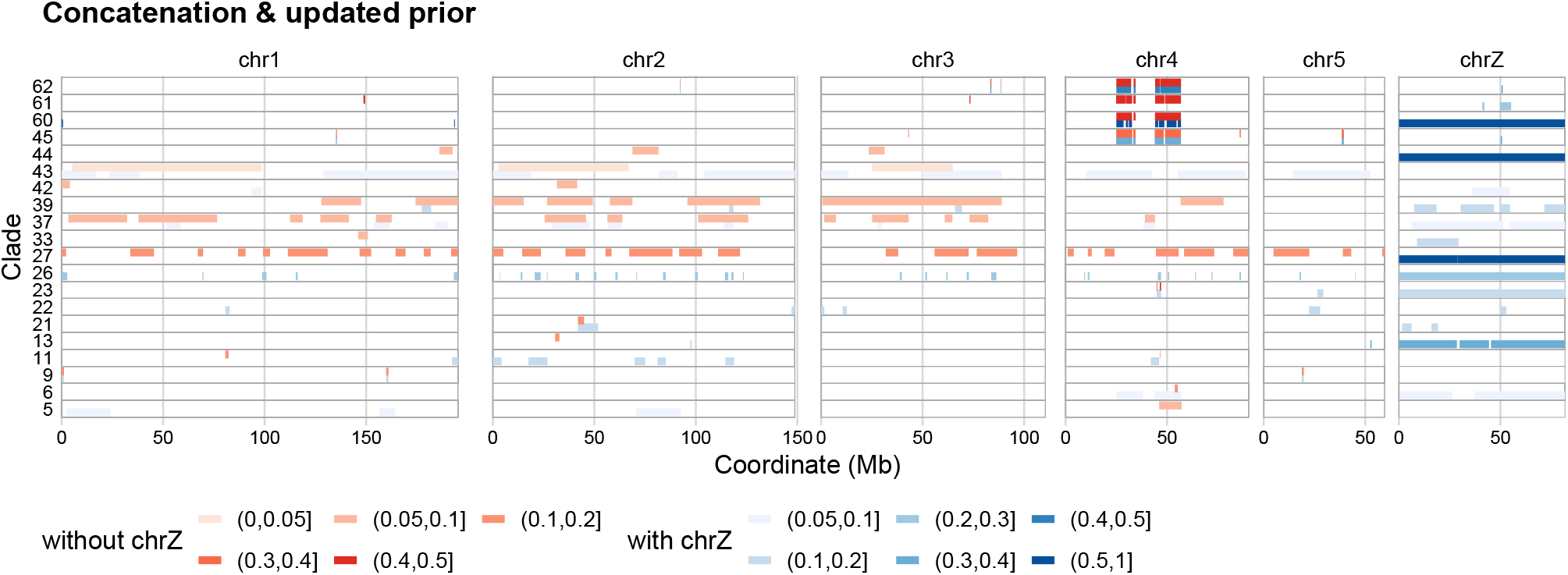
Phlag analysis of bird macrochromosomes and the Z chromosome. We ran Phlag twice, once with and once without chromosome Z included. Analyses that exclude the Z chromosome are shown with red colors (upper part of each row) and those that include the Z chromosome are shown with blue colors (lower part of each row). The analyses that exclude the Z chromosome are identical to those in Fig. 4. In general, inclusion of the Z chromosome reduced the number of autosomal outliers detected by Phlag, although there are a few exceptions where short autosomal regions were detected when the Z chromosome was included (e.g., branch 11). This pattern can be explained by the observation that our model includes only one alternative state; thus, if the Z chromosomes show a stronger pattern of disagreement with the default MSC model, it can prevent the detection of more subtle deviations. Employing iterations is a potential solution.

**Fig. S5.**
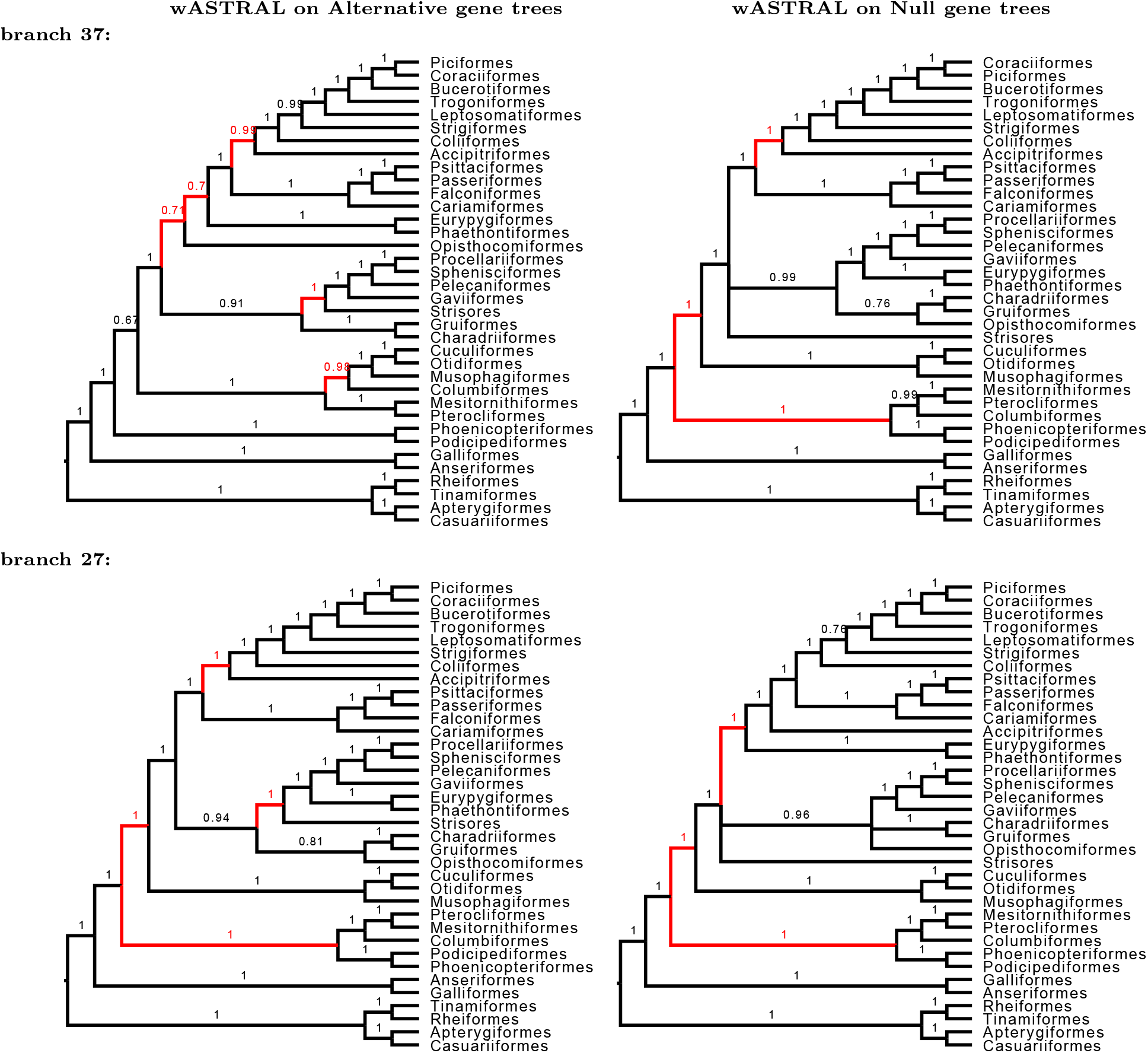
Trees generated by running wASTRAL on gene trees from the null or alternative regions for branches 37 and 27 of birds. This figure shows all avian orders; the versions in Fig. 4 are reduced to show the focal taxa predicted to change. Note that the species tree includes 363 species; we collapse all the orders in the visualizations. We removed orders outside of Neoaves, which we do not study here.

**Fig. S6.**
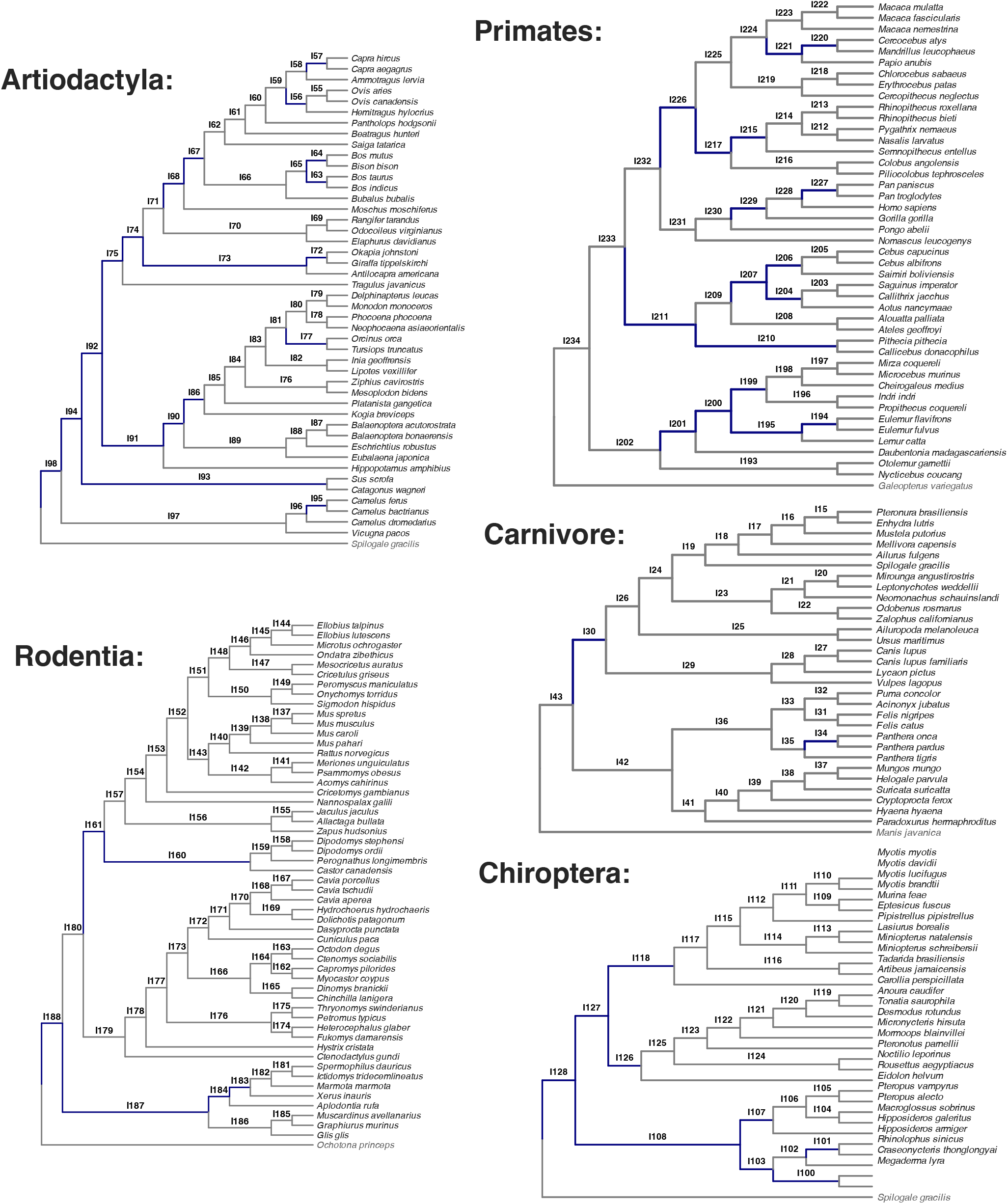
Showing the subtrees from the five key orders selected from the mammalian phylogeny, namely Artiodactyla, Primates, Rodentia, Carnivora, and Chiroptera. Shown clades correspond to the common ancestor node above the edges with regions identified by Phlag with >0.25 Hellinger distance, highlighted in blue.

**Fig. S7.**
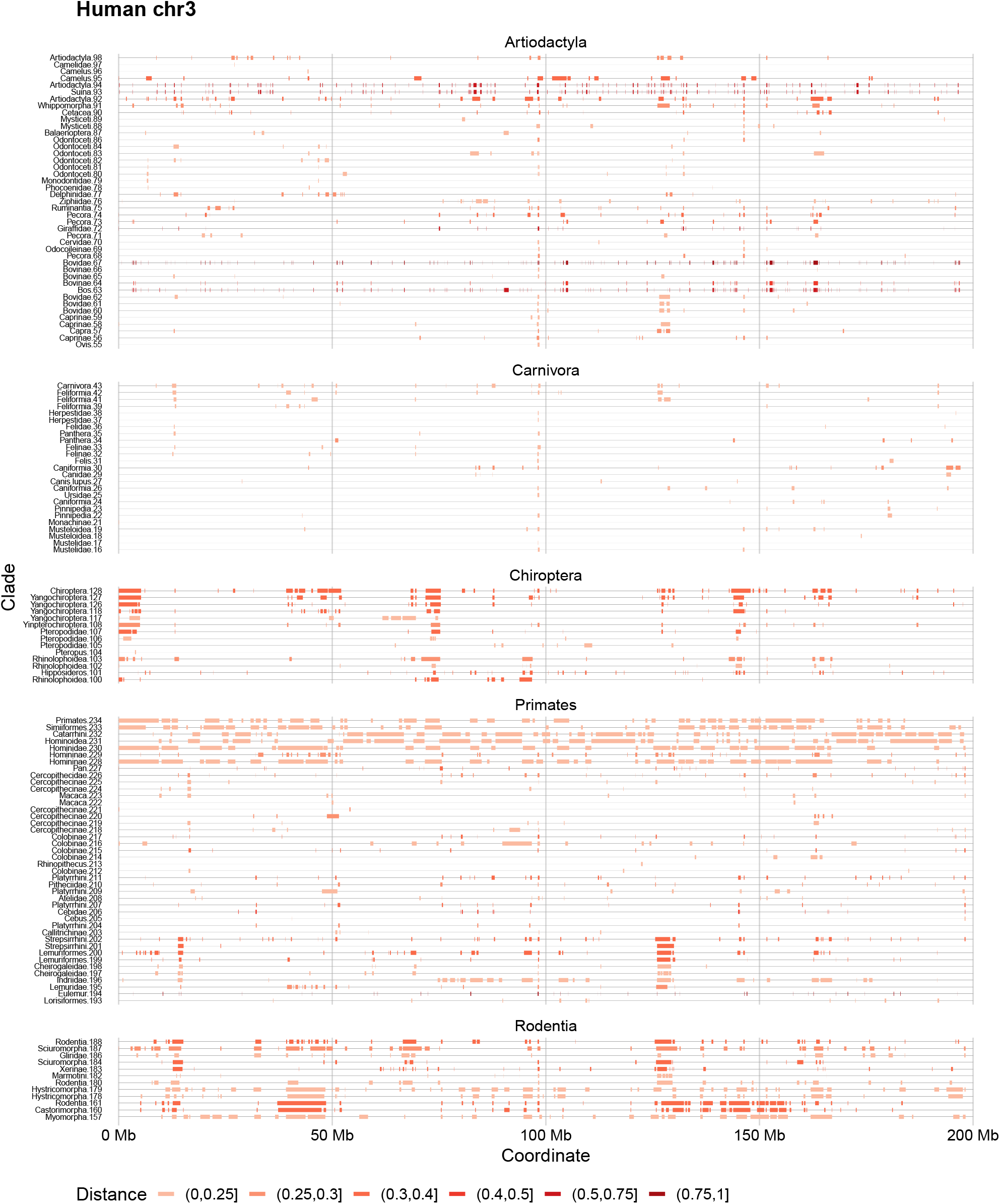
The complete results of the Phlag analysis on the mammal dataset were obtained using the same hyperparameters as the analysis conducted in the manuscript (*β* = 100 and *ρ* = 0.95). Each row corresponds to a branch in the mammal tree. This figure presents results for all individual branches from Carnivora, Chiroptera, Primates, Artiodactyla, and Rodentia. All branches with sufficiently complete data (>90%) are included. In Figure 5, we highlight branches with distances exceeding 0.25. For each branch, we report the lowest common ancestor (LCA) of all taxa in the subtree rooted at the descendant node, along with the order in which this node is visited during the post-order traversal on the *y*-axis.

**Fig. S8.**
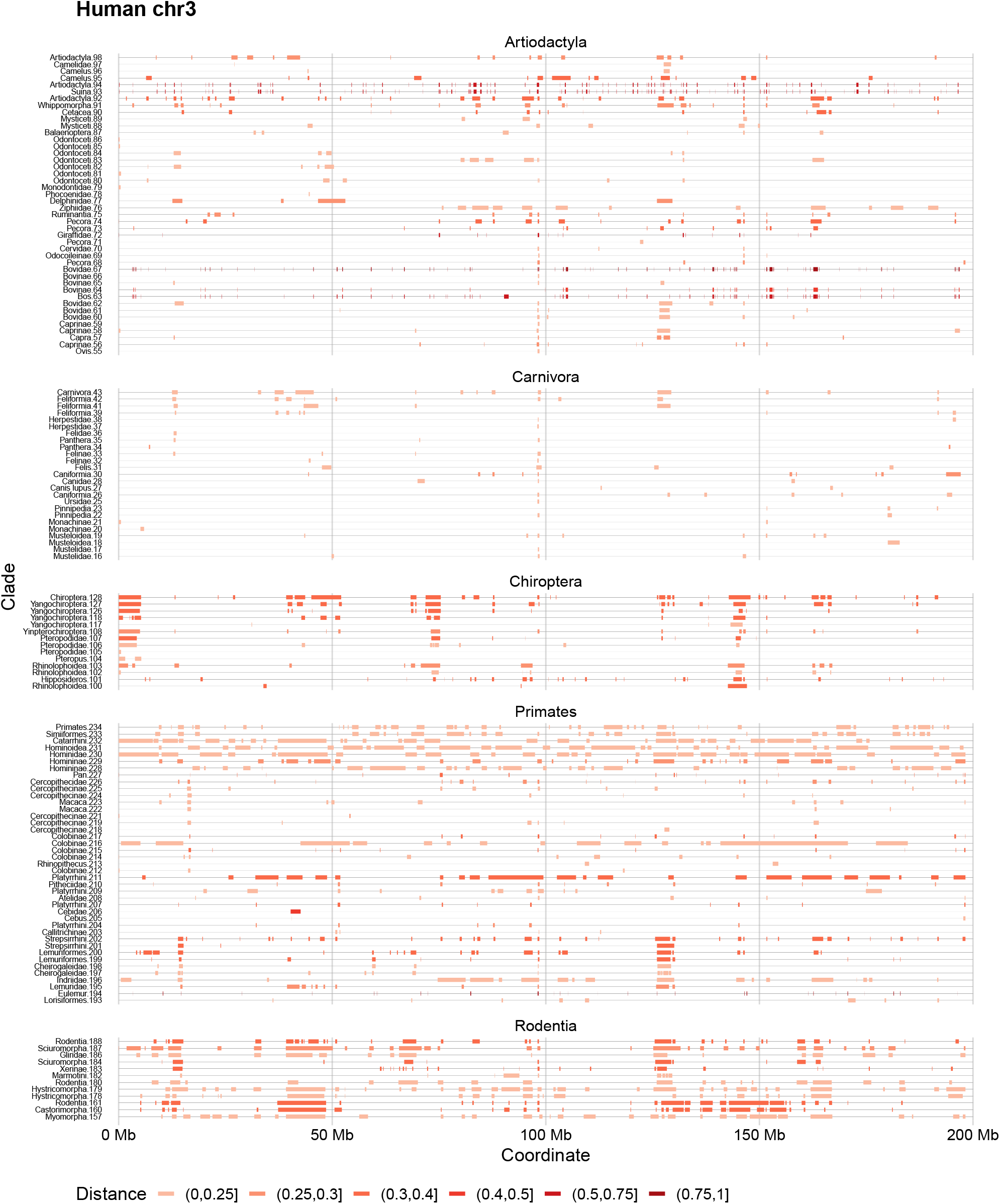
The complete results of the Phlag analysis on the mammal dataset were obtained by setting *β* = 50 and *ρ* = 0.95 to maintain an expected number of transitions per gene tree of 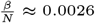, where *N* = 19, 465.

**Fig. S9.**
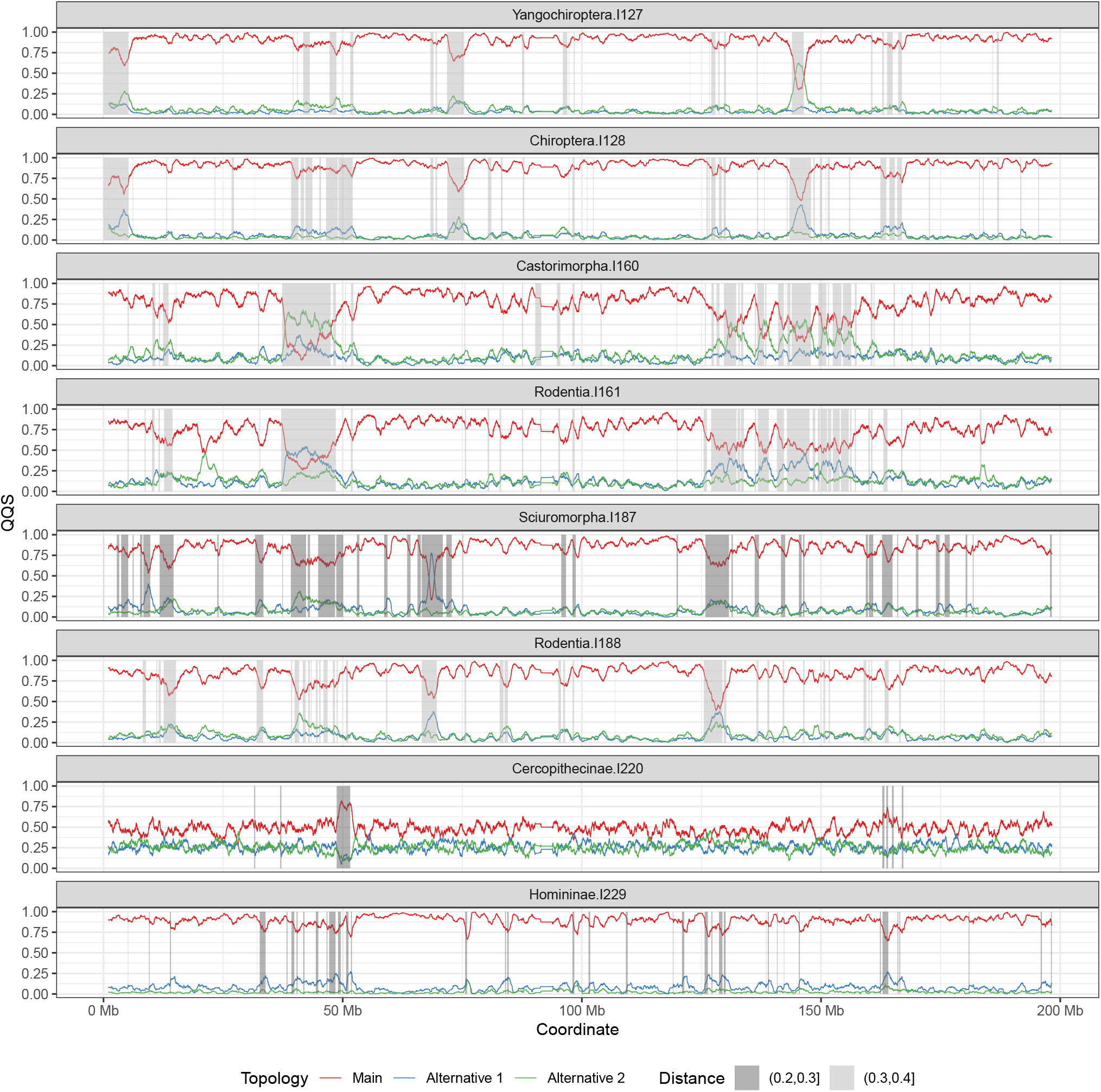
Moving averages of Quartet Quadripartition Support (QQS) values of three topologies for 8 focal branches from Figure S7. Regions identified by Phlag are highlighted in gray, with shading based on the Hellinger distance between the emission distributions of the two states.

## S2. Supplementary Notes

### S2.1. Derivation of the transition matrix priors

We consider a two-state transition matrix

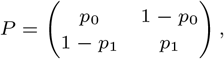

where *p*_0_ = *p*(*Z*_*i*_ = 0 | *Z*_*i*−1_ = 0) and *p*_1_ = *p*(*Z*_*i*_ = 1 | *Z*_*i*−1_ = 1). The stationary distribution *π*^∗^ = [*ρ*,1−*ρ*] satisfies *π*^∗^*P* = *π*^∗^. Solving for *ρ* results in

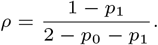

where *ρ* is the prior probability of being in the 0 state.

Let *N* denote the sequence length and let *β* denote the expected number of subsequences with *Z*_*i*_ = 1. Under stationarity, the expected number of 0 → 1 transitions is *Nρ*(1 − *p*_0_). Thus, *Nρ*(1 − *p*_0_) = *β* yields *p*_0_ = 1 − *β* / (*Nρ*). Therefore, we end up with

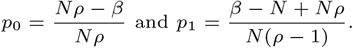

We place conjugate priors *p*_0_ ∼ Beta(*α*_0_, *β*) and *p*_1_ ∼ Beta(*α*_1_, *β*), with expectations 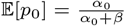 and 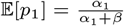. Enforcing the above constraints together with *α*_0_ + *β* = *Nρ* and *α*_1_ + *β* = *N* (1 − *ρ*) gives

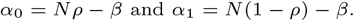

### S2.2. M-step

In the M-step, we update model parameters by maximizing the expected complete log-posterior. Specifically, we solve

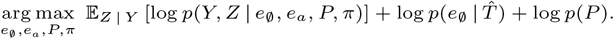

The emission parameters *e*_∅_ and *e*_*a*_ are updated independently across branches by maximizing

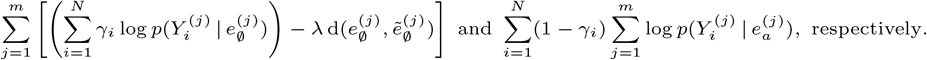

The resulting updates correspond to maximum a posteriori estimates for *e*_∅_ and maximum likelihood estimates for *e*_*a*_. For instance, when 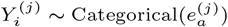, which is the case for the default topology order emissions, MLE is simply

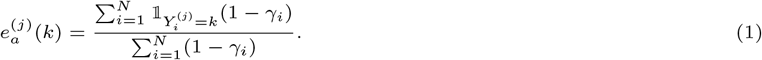

Unlike the maximum likelihood objective of *e*_*a*_, the objective for *e*_∅_ generally admits no closed-form solution due to the distance-based prior term Eq. (3), and hence, the maximization is performed numerically. In particular, we use L-BFGS-B without using the explicit derivative.

The transition probabilities are updated using the expected transition counts as

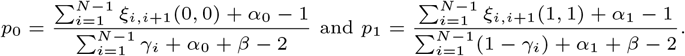

The initial distribution *π* is updated as

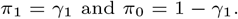

### S2.3. Gene tree simulations

#### Ancestry simulations

We simulate genealogies under the demographic model of the avian species tree inferred by Stiller *et al*. [25] using Hudson’s algorithm implemented in `msprime`. We set the recombination rate to 5 × 10^−9^ and fix the generation time at 10 years across the entire tree. For computational feasibility, we simulate sequences of length 500Kbp and concatenate 12 independent blocks of 500Kbp to obtain 6Mbp-long sequences. We respect these concatenation boundaries when sampling genes for gene tree estimation to minimize the error introduced. We note that the resulting genealogy sequences produced by `msprime` are ultrametric.

#### Incorporating varying rates of evolution

We extend the simulations to incorporate variation in rates of evolution by first scaling the rooted species tree branch lengths, measured in numbers of generations, by branch-specific rate multipliers; i.e., branch lengths in substitution units normalized by the average substitution rate. Note that these lengths are obtained empirically on real data. This produces a non-ultrametric species tree, for which we reassign node times by anchoring the tip with the greatest root-to-tip distance at time zero and shifting all other nodes accordingly. Formally, let *h*(*x*), *d*(*x*), and *t*(*x*) denote the height, depth, and time of a species tree node *x* before scaling, and let *h*′(*x*), *d*′(*x*), and *t*′(*x*) denote the corresponding quantities after scaling. We assign each species tree node a new time *t*′(*x*) = *h*(root) − *d*′(*x*). Since each node in a simulated genealogy belongs to a population corresponding to a species tree node, a genealogy node *y* sampled at time *t*(*y*) within population *x* is assigned a scaled time

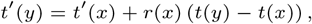

where *r*(*x*) is the rate multiplier of the branch connecting *x* to its parent in the rooted species tree. The effect of the procedure described is to assign a substitution rate to each branch of the species tree and scale each branch of the gene tree by the weighted average of those rates, where the weights are driven by how much time a gene tree branch spends in each species tree branch.

#### Alignment simulations

On these scaled genealogies, we simulate mutations under the GTR model with a mean substitution rate of 1.909 per bp per generation, calculated based on the avian species tree in substitution units. We introduce mutation-rate heterogeneity both across genomic regions and across sites. Specifically, *i*) we partition the genome into random contiguous segments with a mean length of 10^4^ bp and scale the substitution rate of each segment by a random multiplier drawn from a high-variance Gamma distribution. *ii*) Within each segment, we further scale the mutation rate at each site using a Gamma distribution with 20-fold lower variance. The (inverse) variance parameter itself is drawn from a log-normal distribution independently for each independent block.

#### Gene tree estimations

We sample 500bp genes from each 4Kbp window, respecting the concatenation boundaries, and infer gene trees that we use in our benchmarking using IQTREE (v2.4.0) and GTR+G4 model with `--abayes` option enabled.

